# Large scale spatio-temporal study on European hake depicts sensitivity to sea bottom temperature, sea bottom oxygen concentration and sea surface temperature

**DOI:** 10.1101/2023.09.16.558035

**Authors:** Iosu Paradinas, Guillem Chust, Dorleta Garcia, Leire Ibaibarriaga

## Abstract

European hake (*Merluccius merluccius*) is a commercially important fish species that is known to have a marked bathymetric preference. No other environmental variable has yet been identified to drive the distribution of hake. This study looked into different climatic variables at different depths and identified sea bottom temperature, sea bottom dissolved oxygen concentration and chlorophyll concentration to affect the distribution of both juvenile and adult hake distributions.

## 1 Introduction

European hake (*Merluccius merluccius*) is a very important fish species, whose economic importance in the European Union (EU) is among the top 15 species caught and top 10 species consumed. This is especially true for France, Spain, and the United Kingdom, whose fleets catch the most European hake by volume. ıt is found in the Mediterranean and Northeast Atlantic Ocean, where it extends from the north-west African coast to Norwegian Sea waters [8]. Over the last decade, the species has become more common in its northern border, particularly in bottom trawl catches [1, 2, 12].

European hake are carnivorous and primarily feed on small fish, crustaceans, and cephalopods [3]. European hake is well adapted to a wide range of water temperatures and depths, making it a versatile predator in various marine environments.

Reproduction is characterized by the production of large, buoyant eggs and external fertilization. Larvae are pelagic and gradually develop into juvenile fish before settling to the seafloor as adults.

European hake recruits are known to have a marked bathymetric preference to the 150-200m bathymetric strata [7, 9], while adults show a broader bathymetric preference up to 350m deep [10]. However, after a literature review under the SeaWise project and to the best knowledge of the authors, no clear environmental relationships had been found prior to this study, probably due to the restricted spatial scale of most studies. Few studies have mentioned temperature and salinity in their hake studies, but relationships were either vague [5] or not statistically tested.

The objective of this study was therefore to identify environmental variables that characterise the distribution of juvenile and adult European hake. This way we expect to improve our understating on this highly valuable commercial species and improve its future management.

## 2 Materials and Methods

### 2.1 European hake data

We downloaded European hake data from DATRAS covering the North Eastern Atlantic ranging from Spain to Norway, including the BTS, EVHOE, FR-CGFS, ıE-ıGFS, NS-ıBTS, SCOWCGFS, SP-ARSA, SP-NORTH and SP-PORC surveys as shown in Figure 1. The seasonal coverage of each survey was different (Table 1), but given that our aim was to infer climatic relationships (not including any spatial or spatio-temporal latent fields), we did not worry about the temporal mismatch between surveys. We separated European hake in juveniles and adults by considering as Juveniles all those specimens up to 42cm in length, and adults all individuals larger than 42cm. Hake juvenile and adults’ data was then transformed to presence-absence data to fit logistic models.

**Table 1.**
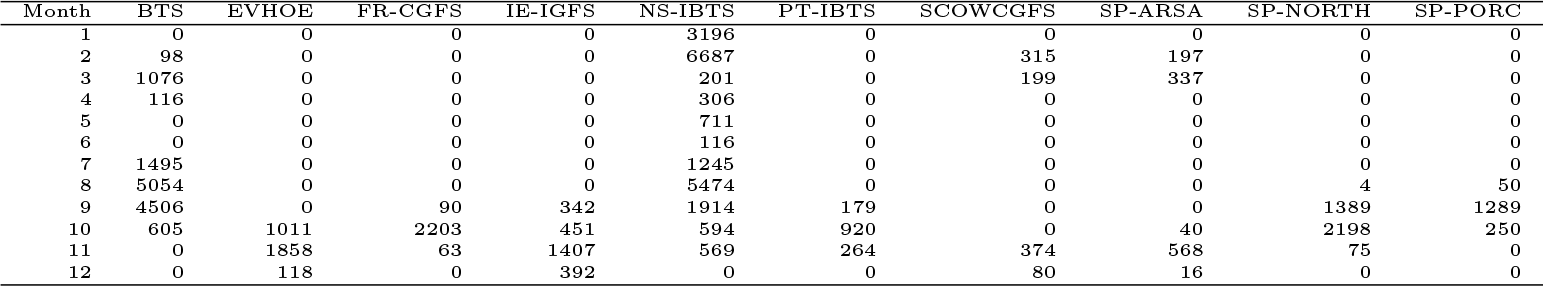
Surveys seasonal distribution. Values are the number of samples collected by each survey between 1993 and 2021.

**Figure 1.**
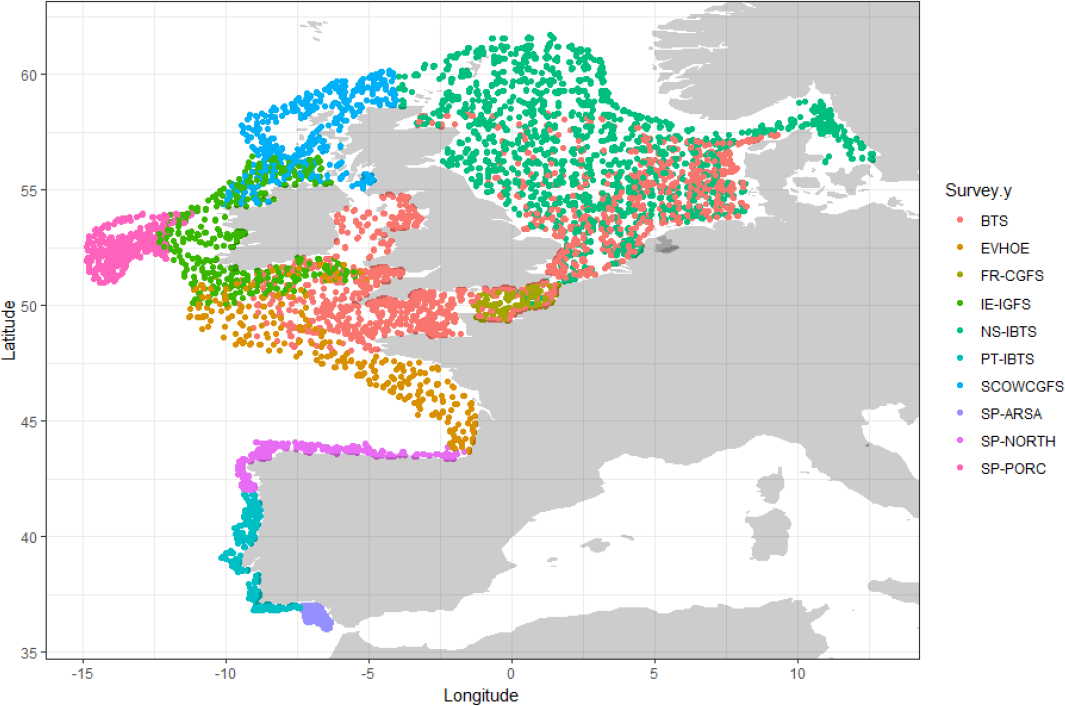
Trawl surveys’ spatial distribution

### 2.2 Environmental and climate projection variables

We used climate data products developed through the FutureMares project. These data are available at zenodo.org and provide both historical and projection data on temperature, salinity, dissolved oxygen concentration, ph and chlorophyll concentration at the sea surface, at 25 meters of depth and at the sea bottom (with the exception of chlorophyll). Historical climate data come from the GLORYS12V1 reanalysis provided by the Copernicus Marine Environment Monitoring Service that was used to train projection models. Future climate projections combine Shared Socioeconomic Pathway (SSPs) inputs and Representative Concentration Pathways (RCPs) inputs and ensembled different projection models for three different scenarios: SSP1-RCP2.6, SSP2-RCP4.5, and SSP5-RCP8.5.

### 2.3 Modeling

Given the large number of explanatory variables, we first performed a broad variable selection process combining three different approaches. First we fitted several univariate splines using generalised additive models (GAM), quadratic effects using generalised linear models (GLM) and Shape-Constrained Generalised Additive Models (SC-GAMs) to visually check that fitted relationships in all three methods were consistently in line with the ecological niche theory [6] inferring unimodal or concave relationships. We filtered out those explanatory variables that showed inconsistent process-environmenta relationships. Then, we applied two Machine Learning (ML) algorithms, random forest and Bayesian additive regression trees, over the remaining variables to further select those climatic variables that showed the biggest impact in predicting hake juveniles and adults’ distribution.

The remaining environmental variables were modelled using SC-GAMs after checking that variables did not show high correlation coefficients. SC-GAMs are based on the same statistical framework as GAMs and we used them due to their capacity to constrain the functional response curve of the fitted effects to shapes that are in agreement with the niche theory [4]. Despite bathymetry being a well-known environmental driver for hake [7, 9, 10], it could also mask the effect of other more dynamic environmental variables, thus was left behind in the model selection process.

The final variable selection process was performed using two different approaches. On the one hand, we performed the forward stepwise variable selection based on AıC using a cut-off value of two AıC scores. On the other hand, we used a ten-fold cross validation (CV) approach to do the forward stepwise selection based on its predictive capacity. Each cross validation used 80% of the data to train the model and 20% to check the predictive capacity of the model. We finally chose CV based results as they were more restrictive. The whole process was performed both for hake juveniles and adults.

## 3 Results

After the first variable selection process using visual validation and ML algorithms, sea bottom temperature, sea bottom dissolved oxygen concentration, salinity at 25m and chlorophyll concentration at the surface were kept as potential explanatory variables to be included in our SC-GAM models. Final SC-GAM models dropped salinity in both hake juvenile and adults (Figure 2).

**Figure 2.**
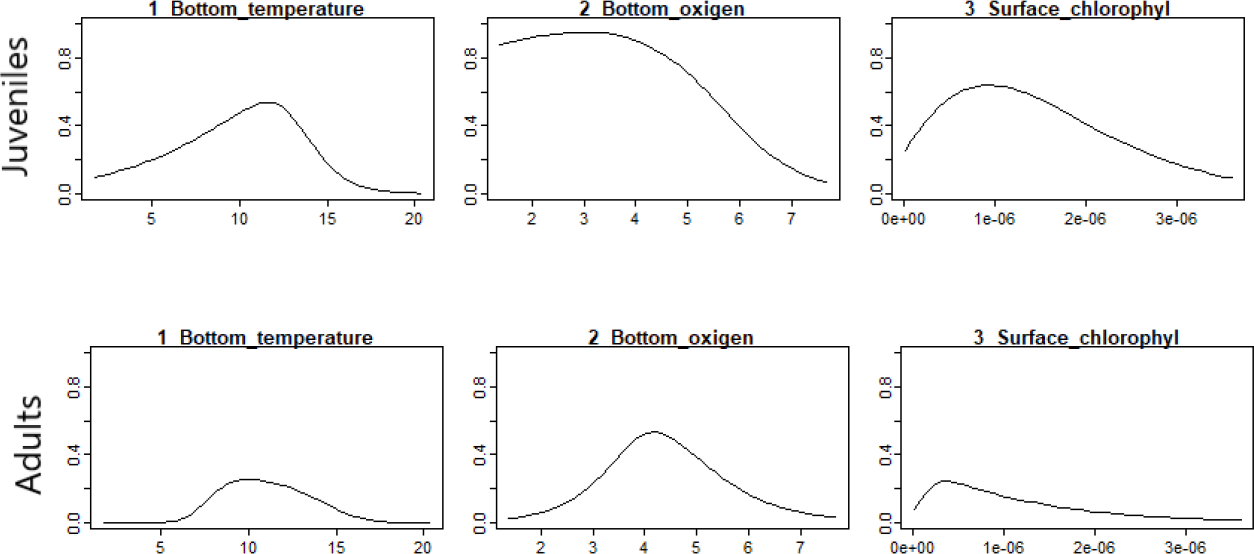
Fitted environmental effects in the hake juveniles (top) and adults (bottom). Each plot displays the marginal functional response of each environmental variable with regards to occurrence of hake.

Sea bottom temperature showed a maximum hake adult and juvenile occurrence around the ten degree Celsius mark (left panels in Figure 2). Hake juvenile’s maxima was at slightly higher temperatures than the adults one, which is in accordance with their bathymetric preferences where adults prefer deeper waters that would generally be colder. Regarding sea bottom oxygen concentration, both life stages seem to avoid high values (central panels in Figure 2). Adults seem to avoid low oxygen concentrations while juveniles seem to do well there. Lastly, highest hake occurrence seem to occur at relatively low chlorophyll concentration areas (right panels in Figure 2).

Based on these models we also produced future projections for hake juvenile and adult populations for every month of every decade until 2099 using SSP1-RCP2.6, SSP2-RCP4.5 and SSP5-RCP8.5 scenario projections. Annex 4 shows a summary of these projections for the month of october.

## 4 Discussion

European hake (*Merluccius merluccius*) is a very important fish species whose distribution has long been studied. This study looked into the climatic variables that drive its large scale distribution. To do so we have used hake occurrence data covering the North Eastern Atlantic from Spain to Norway (Figure 1). Results identified that both juveniles and adults preferred sea bottom temperatures around 10 degrees Celsius and seemed to avoid high chlorophyll and high sea bottom oxygen concentrations.

We tested several climatic variables: temperature; salinity; dissolved oxygen concentration; ph; and chlorophyll concentration. Each of these variables were extracted at different depths: surface; 25m deep; and at the sea bottom. Random forest and Bayesian additive regression trees highlighted the importance of sea bottom temperature, sea bottom dissolved oxygen concentration, chlorophyll concentration at the surface and salinity at 25m. Finally, salinity was dropped from the final models using SC-GAMs but could probably be tested again in the future with more data.

Future European hake distribution studies could use these results as valuable prior knowledge to fit their models. All data used in this study and the script used to fit the final models are available at XXXGithublink. This way one could combine their newly collected data with the data used in this study to get better informed models. Given that the data used in this study was occurrence data, if new hake distribution studies use other types of data (e.g. abundance or biomass), integrated species distribution models may provide a good statistical framwork to combine the datasets into a single model [11].

## Acknowledgments

We thank all those who took part in the literature review within the Seawise project as well as all those who put their effort in conducting the trawl surveys used in the study.

## Annex

### Future hake distribution projections

**Figure 3.**
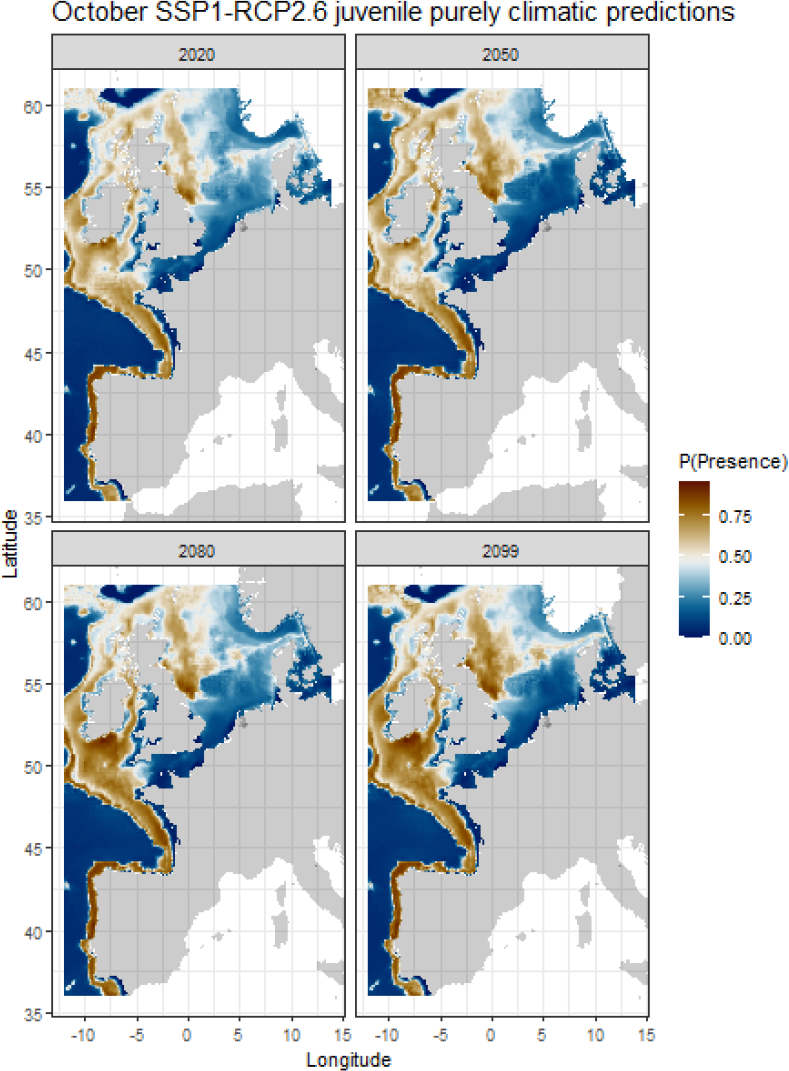
Climatic model predictions for juvenile hake in October for the years 2020, 2050, 2080 and 2099 under the SSP1-RCP2.6 climatic scenario.

**Figure 4.**
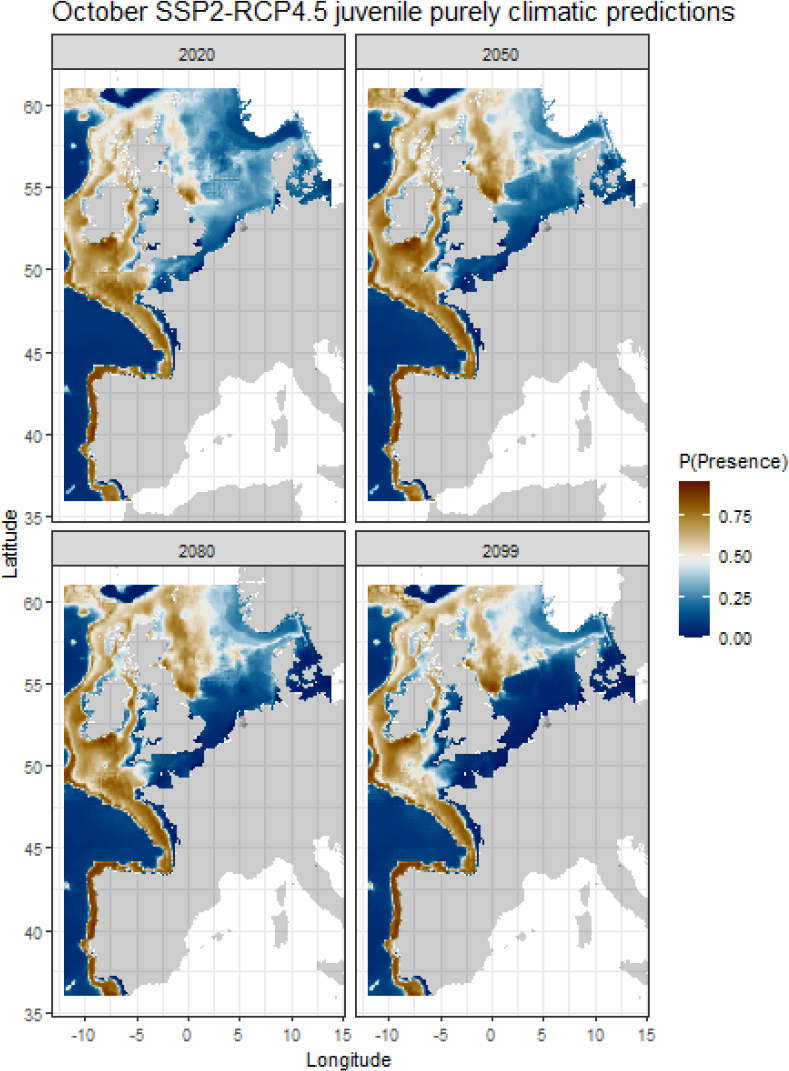
Climatic model predictions for juvenile hake in October for the years 2020, 2050, 2080 and 2099 under the SSP2-RCP4.5 climatic scenario.

**Figure 5.**
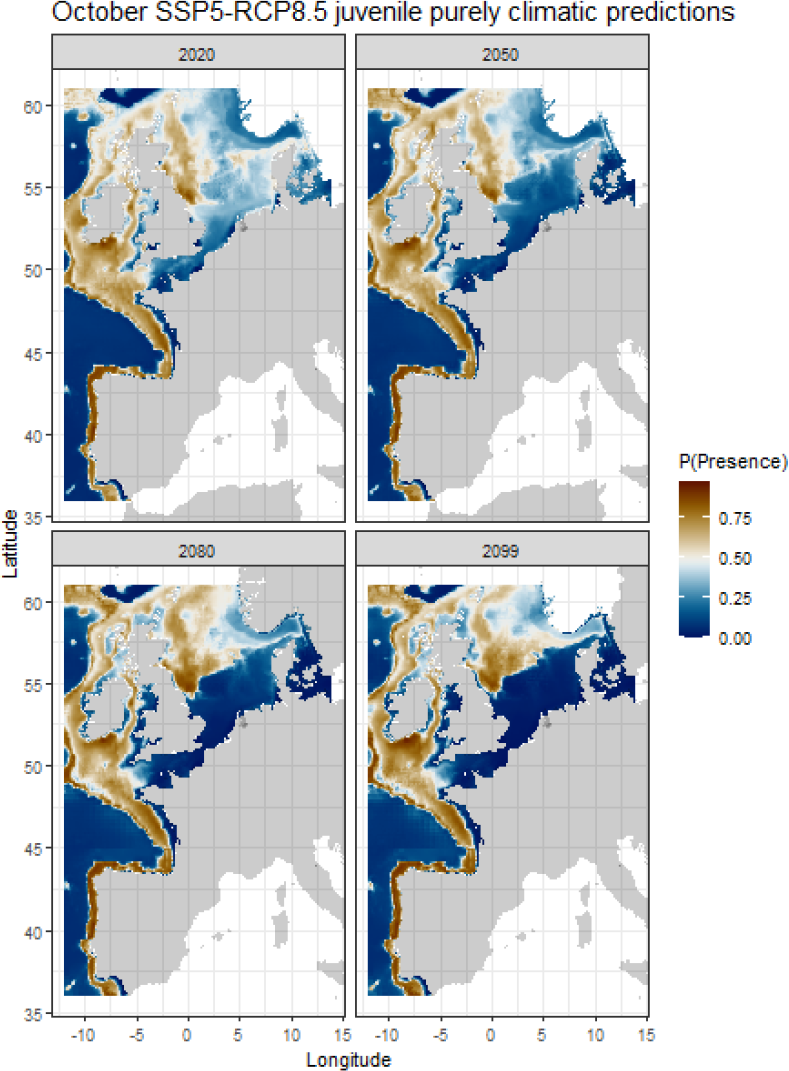
Climatic model predictions for juvenile hake in October for the years 2020, 2050, 2080 and 2099 under the SSP5-RCP8.5 climatic scenario.

**Figure 6.**
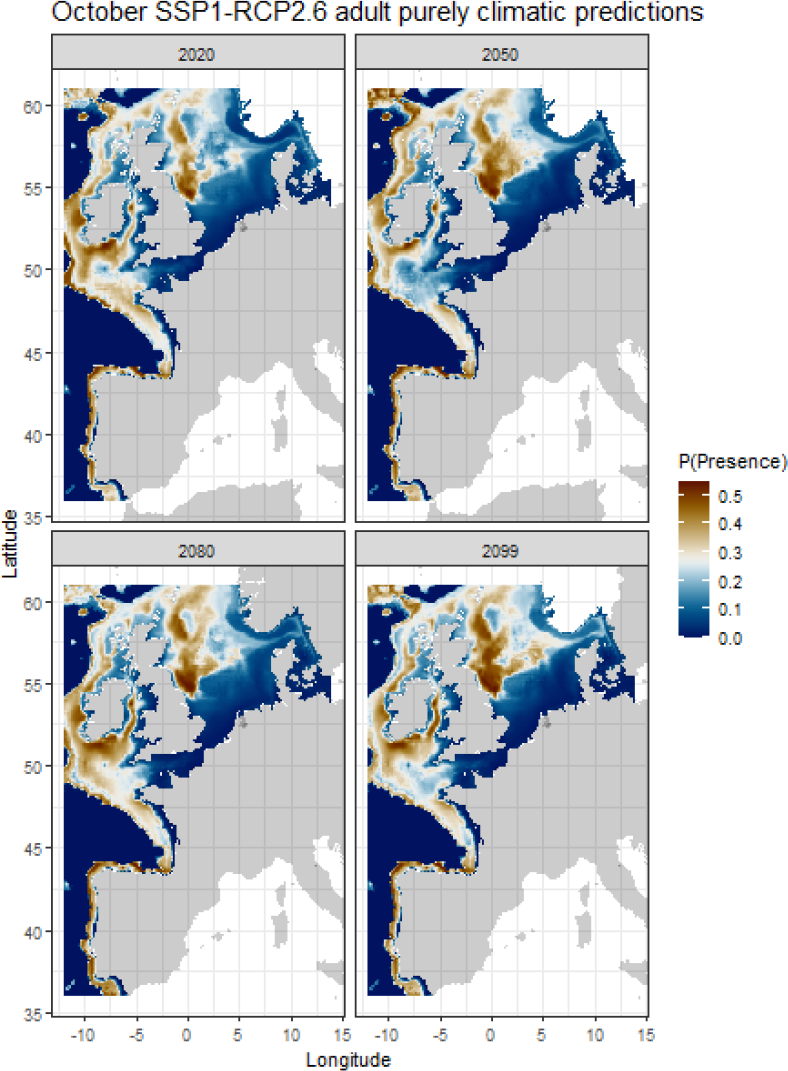
Climatic model predictions for adult hake in October for the years 2020, 2050, 2080 and 2099 under the SSP1-RCP2.6 climatic scenario.

**Figure 7.**
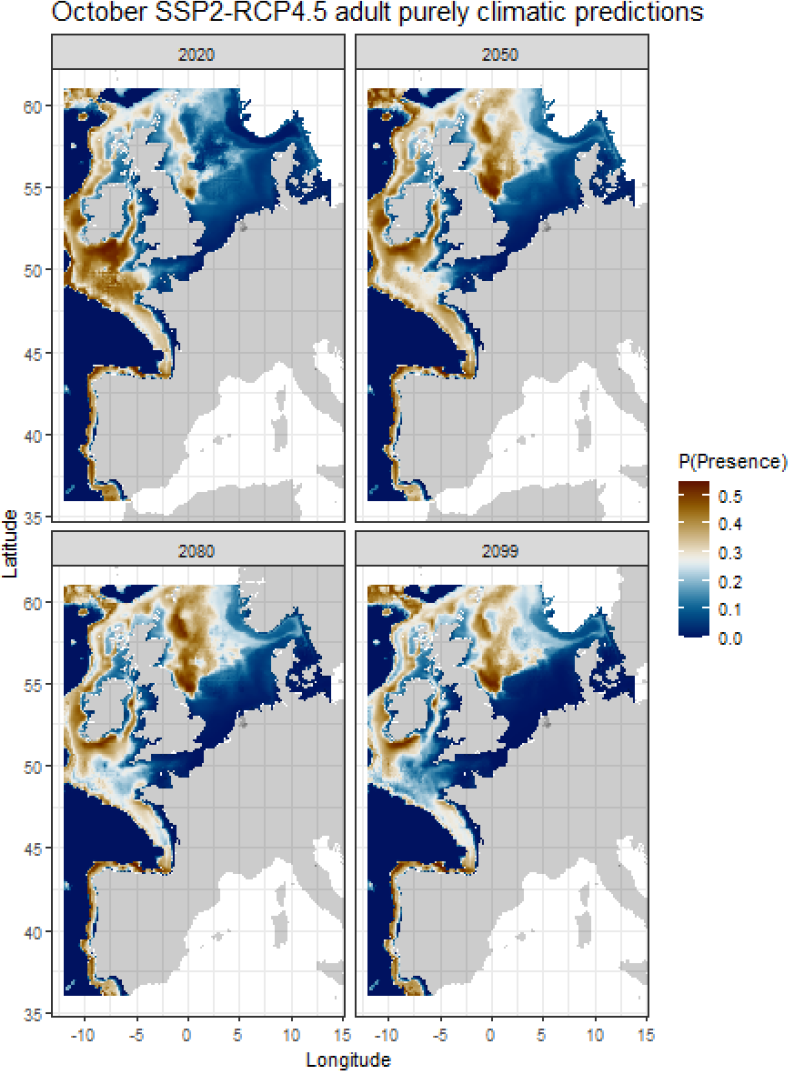
Climatic model predictions for adult hake in October for the years 2020, 2050, 2080 and 2099 under the SSP2-RCP4.5 climatic scenario.

**Figure 8.**
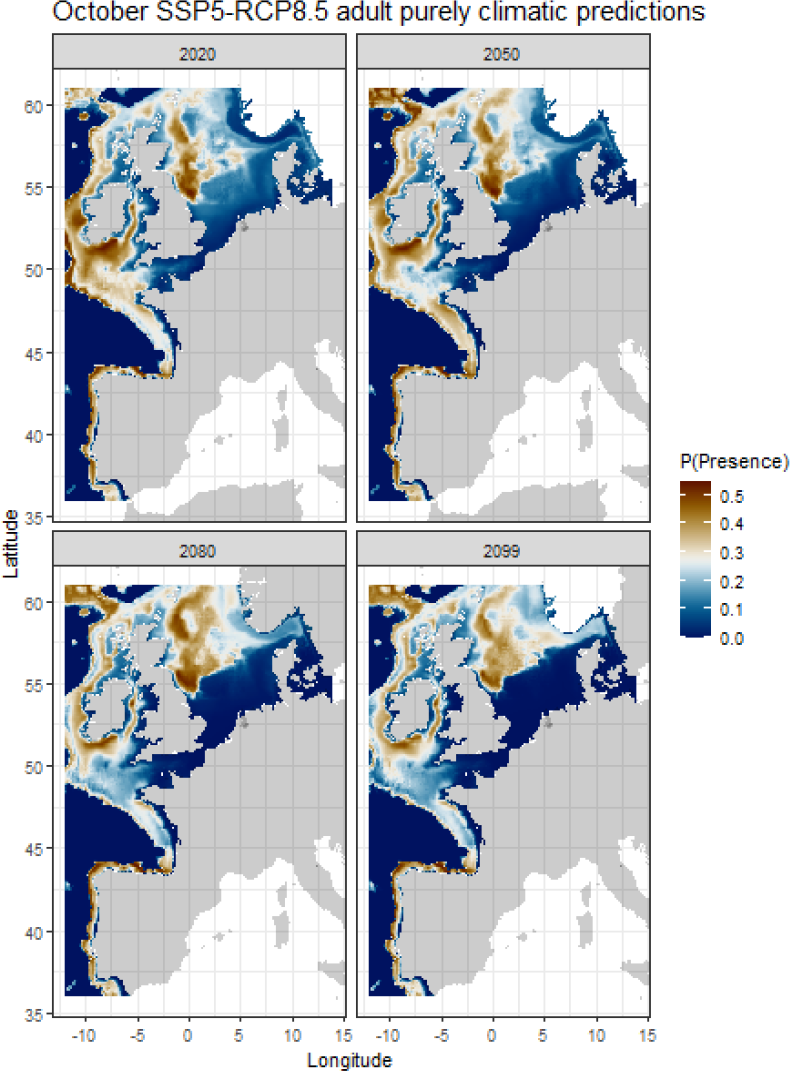
Climatic model predictions for adult hake in October for the years 2020, 2050, 2080 and 2099 under the SSP5-RCP8.5 climatic scenario.

